# Versatile Learning without Synaptic Plasticity in a Spiking Neural Network

**DOI:** 10.64898/2026.01.01.697291

**Authors:** Kai Mason, Navid Akbari, Aaron Gruber, Wilten Nicola

**Affiliations:** Department of Cell Biology and Anatomy, Cumming School of Medicine, University of Calgary, Calgary, Alberta, Canada; Hotchkiss Brain Institute, University of Calgary, Calgary, Alberta, Canada; Department of Physics and Astronomy, Faculty of Science, University of Calgary, Calgary, Alberta, Canada

## Abstract

Learning in the brain’s cerebral cortex is widely attributed to synaptic plasticity among cortical neurons. However, a growing body of evidence suggests that alternative processes, such as modulation of intrinsic excitability or gating by subcortical inputs, may also serve as important learning mechanisms. We developed the Bias Adaptive Neural Firing Framework (BANFF) as a simplified model of such phenomena embodied by a learnable bias current for each neuron of a rate-based network. Here, we extend this framework to spiking neural networks. We show that learning such biases enables one recurrent spiking neural network with fixed and random synaptic weights to perform well on nine tasks spanning classification, regression, and closed-loop dynamical systems mimicry. The network learnt a unique bias set for each task, and unlike recurrent synapse-based learning, new learning did not interfere with previous learning. The network was robust to non-stationary F-I curves (spike frequency adaptation), and biases could be learned with a learning algorithm (e-prop) that is more biologically plausible than stock gradient descent. Overall, we show that the BANFF can be extended from rate-based to spiking neural networks, maintaining good multi-task performance with a single network of spiking neurons.

## Introduction

A prevailing hypothesis is that learning and memory in cerebral neural circuits arise primarily through changes in synaptic strengths amongst cortical pyramidal neurons. This view is supported by decades of experimental and theoretical work on synaptic plasticity, from longterm potentiation to spike-timing-dependent plasticity (STDP) [1]. Consequently, most computational models of neural networks place synaptic plasticity as integral to learning [2]. Yet, neurons possess other adjustable biophysical parameters, and growing evidence indicates that intrinsic excitability can also be modified on behaviourally relevant timescales [3].

Activity and neuromodulators can shift excitability by altering ion channel expression, resting potential, spike threshold, or leak conductances, with changes that persist beyond the inducing stimulus [3–8]. Such activity modulations effectively bias a neuron towards or away from spiking, reshaping its input–output curve without altering its synaptic couplings. At the population level, subcortical inputs can shift bias currents across many cortical neurons simultaneously, providing a systems-level handle on circuit dynamics and state [9–13]. These observations suggest a complementary route to learning in which synaptic couplings remain largely fixed while biases act as control parameters for network dynamics.

Research in machine learning has investigated how useful computations can emerge from networks without changing recurrent synapses. Reservoir computing showed that random recurrent networks paired with adaptive readouts can solve complex tasks [14, 15]. More recent work has examined learning via modulation of neuronal excitability, gain, or bias terms, demonstrating that rich dynamics and task acquisition are possible with fixed recurrent connectivity [16–18]. Building on this theme, our recent Bias Adaptive Neural Firing Framework (BANFF) established that feedforward or recurrent neural networks with random and fixed synaptic connectivity can nevertheless support rapidly reconfigurable dynamic computing when only bias-like terms are adapted [19]. This rate-based network model achieved fast task switching without altering recurrent weights, and highlighted low-rank structure as a parsimonious scaffold for controllable dynamics.

Here we extend the BANFF along biophysical and algorithmic axes by moving from rate-based dynamics to a spiking formulation with adaptive leaky integrate-and-fire (ALIF) neurons [20]. In this setting, synapses remain fixed and constrained (low-rank, distinct excitatory and inhibitory populations), whilst learning acts exclusively through per-neuron biases. The adaptive leaky integrate and fire (ALIF) model renders the neuron’s *F* − *I* curve non-stationary and history-dependent, so that a ‘bias’ is not merely a static offset but a state- and time-dependent control variable. We use e-prop–style eligibility traces and surrogate gradients [20] to obtain a more biologically plausible learning algorithm based on local three-factor updates for temporal credit assignment. We found that bias-only learning is sufficient to tune network trajectories towards complex temporal behaviours, and switching among biases can control excitability for rapid, state-dependent reconfiguration atop a stable recurrent scaffold, suggesting a plausible mechanism for functional reconfiguration of the brain’s cortical processing.

## Results

### Low-Rank Spiking Neural Networks can be Trained with Bias Currents Only

We considered a single fixed–synapse recurrent spiking neural network (SNN) formulated similarly to our previously described rate-based BANFF networks (Figure 1A-B). The network comprised 32,000 ALIF spiking neurons with low–rank recurrent structure [21–26] and with neurons that were either exclusively excitatory or inhibitory (i.e., satisfied Dale’s Law) (Figure 1B) [27]. To learn the bias currents for each neuron, we utilised e-prop [20], a more biologically plausible gradient-based algorithm than backpropagation through time, which uses eligibility traces to solve the temporal credit assignment problem during training. Learning acted only on per–neuron biases via e–prop updates with a fixed broadcast decoder [20]; the recurrent and input/output synapses were held fixed throughout, and all biases were initialised to the spike threshold (−50 mV). Two linear decoders were present: the primary head that shared the recurrent SNN’s synaptic filter (double exponential synapse, 2 ms rise and 50 ms decay time); and a downstream secondary readout-head driven by its own fixed two–stage synaptic filter (double exponential synapse, 10 ms rise and 200 ms decay time) with learned weights (no feedback into the core) (Figure 1A). Note that the readout head does not impact the dynamics of the network at all, and is more akin to how an experimentalist would “decode out” the behaviours that the network engages in. Inputs were *z*–scored and scaled by 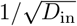 (size–invariant drive).

**Figure 1.**
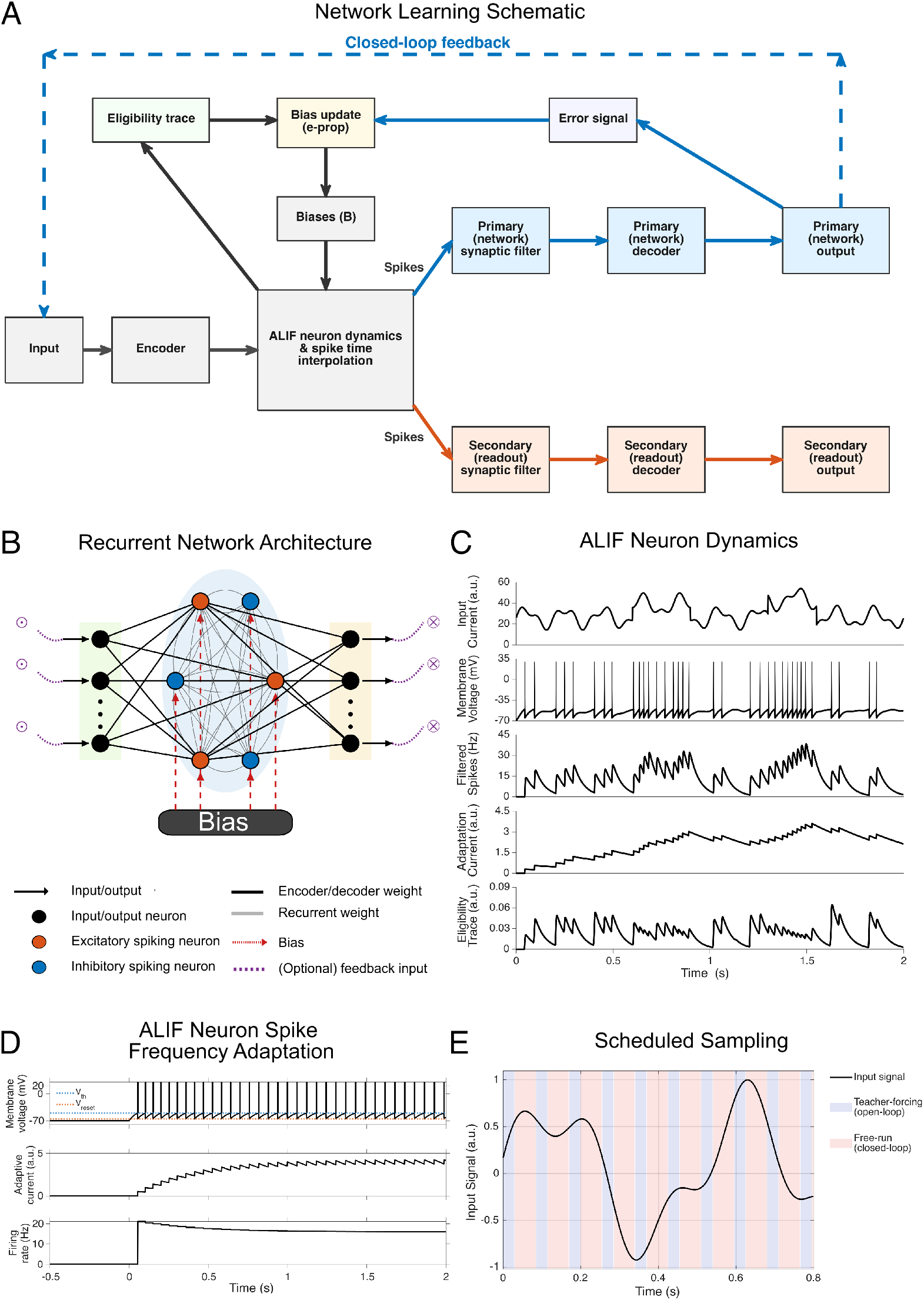
Schematic of Spiking BANFF Networks. (A) A flowchart of the spiking neural network (SNN) training framework used in this study. (B) A graphic representation of the SNN used in this study showing the inputs, recurrent population of spiking neurons, outputs, feedback inputs and trainable biases. The network has been scaled down for graphic purposes and is not a true representation of the number of neurons. (C) Example dynamics of the adaptive leaky integrate and fire (ALIF) neuron showing the input current, corresponding membrane voltage (with cosmetic spikes added for visual clarity), the filtered spikes, the adaptive current and the eligibility trace for a period of 2 s of simulation time. (D) The membrane voltage, adaptive current and firing rate (measured with inter-spike interval) of a single ALIF neuron with a constant input current. Demonstrating the spike-frequency adaptation of the neuron in response to a constant current. (E) The scheduled sampling procedure used in this study shows interleaved periods of open-loop supervisor input and closed-loop inputs.

We first trained the network on three continuous–time systems: two chaotic attractors (Lorenz and Sprott S systems) and one oscillator (Van der Pol Oscillator). Supervision alternated between 30 ms teacher–forcing and 55 ms closed-loop free–run segments with off-policy supervision within each window. Evaluation consisted of (50 s) closed–loop rollouts from five unseen initial conditions per system. The network’s performance was periodically assessed by phase portrait similarity with the true system, and the best-performing set of learned biases and secondary decoder readout weights was selected using this criterion (Methods and Supplementary Material).

Across all three systems, the learnt dynamics reproduced the qualitative geometry of the target (Table 1 and Figure 2): closed–loop phase portraits corresponded to the ground–truth manifolds (Supplementary Figure 1); Poincaré section point clouds aligned in location and spread (Supplementary Figure 2); and return–map pairs preserved the structure of the underlying map (Supplementary Figure 2). As expected for a spiking model with fixed synapses, the learnt portraits were noisier than the ground truth systems in all cases, but were localised to the same region of the portrait. In the chaotic cases, the time-series trajectories diverged (Supplementary Figure 3), but the invariant–set geometry and section statistics corresponded to the ground–truth distributions, consistent with correct identification of attractor geometry rather. For the oscillator, there was a slight phase shift with the true trajectory attributable to a mismatch in the learned frequency (Supplementary Figure 3), which is common in learned oscillators.

**Table 1:** Network performance. Dynamical systems rows report qualitative visual geometry on long, strictly closed–loop rollouts. Classification and regression were evaluated open–loop with 300 ms presentations. All numerical values refer to the secondary readout. See Supplementary Material for full task details.

| Name | Eval | Metrics | Notes |
| --- | --- | --- | --- |
| **Dynamical systems** |  |  |  |
| Lorenz system | closed | **Phase portrait:** Visually similar<br>**Poincaré section:** Visually similar<br>**Return map:** Localised to the same region, similar spread, visibly noisy. | Task to mimic a dynamical system closed-loop. Phase portraits, Poincaré sections and return maps localised to the same region as those of the true systems. Network outputs were noisier than the true systems across all visualisations. Time series diverged for chaotic systems but maintained system-level metrics [44]. |
| Sprott S system | closed | **Phase portrait:** Visually similar<br>**Poincaré section:** Localised to the same region, similar spread, visibly noisy.<br>**Return map:** " " | " " [45] |
| Van der Pol oscillator | closed | **Phase portrait:** Visually similar | " " [46] |
| **Classification** |  |  |  |
| Wisconsin Diagnostic Breast Cancer | open | **Accuracy (%)**: 99 | Task to classify breast mass as malignant or benign based on 30 features [47]. |
| MNIST | open | **Accuracy (%)**: 92 | Task to classify hand-written digits from $28 \times 28$ images (10 classes) [48]. |
| Fashion MNIST | open | **Accuracy (%)**: 84 | Task to classify items of clothing from $28 \times 28$ images (10 classes) [29]. |
| **Regression** |  |  |  |
| Car price | open | **Signed error (mean ± SD)**: £64 ± £1,736<br>**Pearson $r$**: 0.96 | Task to output car price from eight features. True test prices range from £1,200 to £51,000. |
| Abalone age | open | **Signed error (mean ± SD)**: -0.19 ± 2.04 years<br>**Pearson $r$**: 0.78 | Task to predict abalone age based on eight features. True test ages range from 3 to 27 years. |
| Yacht Hydrodynamics | open | **Signed error (mean ± SD)**: **0.35** ± **1.79**<br>**Pearson $r$**: 0.99 | Task to predict residual resistance per unit weight of displacement of yachts based on six features. True test values range from 0.01 to 55.44. |

**Figure 2.**
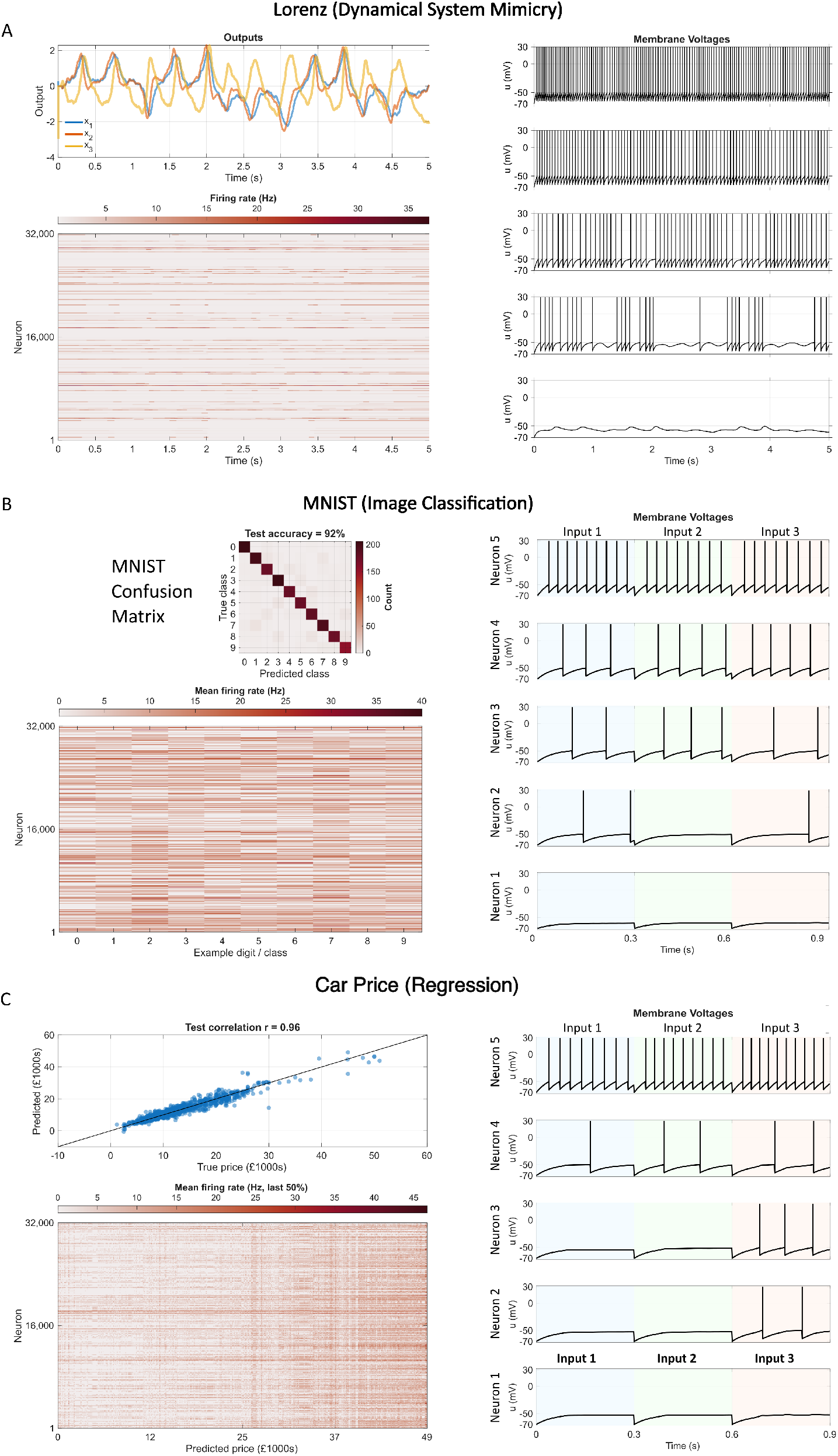
Trained BANFF Networks Switching Between Tasks. (A) The closed-loop network secondary readout output (top left), neural firing rates (bottom left) and five example neuron membrane voltages (right) for the Lorenz system task for five seconds of simulation time. (B) The classification accuracy (top left), example neural firing rates for one of each class of image (bottom left) and example membrane voltages for five neurons for three different input images (right) for the MNIST classification task. (C) A scatter plot of the network and true prices (top left), neural firing rates as a function of the output value (bottom left) and five example neural membrane voltage for three input feature sets (right) for the car price regression task. All firing rates were computed using the inter-spike interval (ISI). The five example neuron membrane voltages are equally sampled from the highest to the lowest average firing rate. Different background colours represent different input feature sets for the classification and regression tasks.

We next evaluated the network’s performance on three open-loop classification tasks, using three datasets: Wisconsin Diagnostic Breast Cancer, MNIST, and Fashion MNIST. The data splits were 60*/*20*/*20 (train/validation/test). Per–sample logits were averaged over the final half of a 300 ms presentation window during which the network inputs were held constant, while we used a cross–entropy loss for training. We selected the biases and secondary readout weights based on the epoch with the highest validation accuracy. The secondary decoder head achieved accuracy comparable to published baselines of similar capacity across all three datasets (Table 1 and Figure 2) [17, 19, 28, 29].

Finally, we tested the network’s performance on three open-loop regression problems using the same simulation mechanics as the classification tasks: car price, abalone age, and yacht hydrodynamic estimation with identical data splits as the open-loop classification task. The targets and predictions were compared using signed mean error ± standard deviation (SD) and Pearson correlation *r*. The network achieved low errors with high correlations on all three problems, comparable to published baselines of a similar capacity (Table 1 and Figure 2) [19, 30, 31].

### Trained BANFF Networks Do Not Use Precise Spike Times

To analyse whether the BANFF network with ALIF neurons learned a rate- or time-encoding representation of the tasks, we conducted perturbation experiments on the closed-loop dynamical systems tasks. For each task, we performed a closed-loop simulation using the learned biases for that task. After the simulation, we jittered spike trains by randomly shifting individual spike times, with each temporal shift drawn from a Gaussian distribution with zero mean and standard deviation *σ* ∈ {0, 5, 13, 32, 80, 200} ms (Figure 3). After jittering, the spikes were fed through the secondary readout matrix and paired synaptic filters (double exponential, 10 ms rise, 200 ms decay times) to extract the time-series. The phase portraits show a gradual and minor decay in fidelity for all systems from *σ* = 0 ms to *σ* = 32 ms and then a larger drop from *σ* = 80 ms and *σ* = 200 ms (Figure 3). These results are more consistent with a rate-encoding scheme than a time-encoding scheme, because a large decrease in the phase portrait quality would be expected for smaller spike-timing jitter amounts for a time-encoding scheme relying on millisecond-level spike times [32].

**Figure 3.**
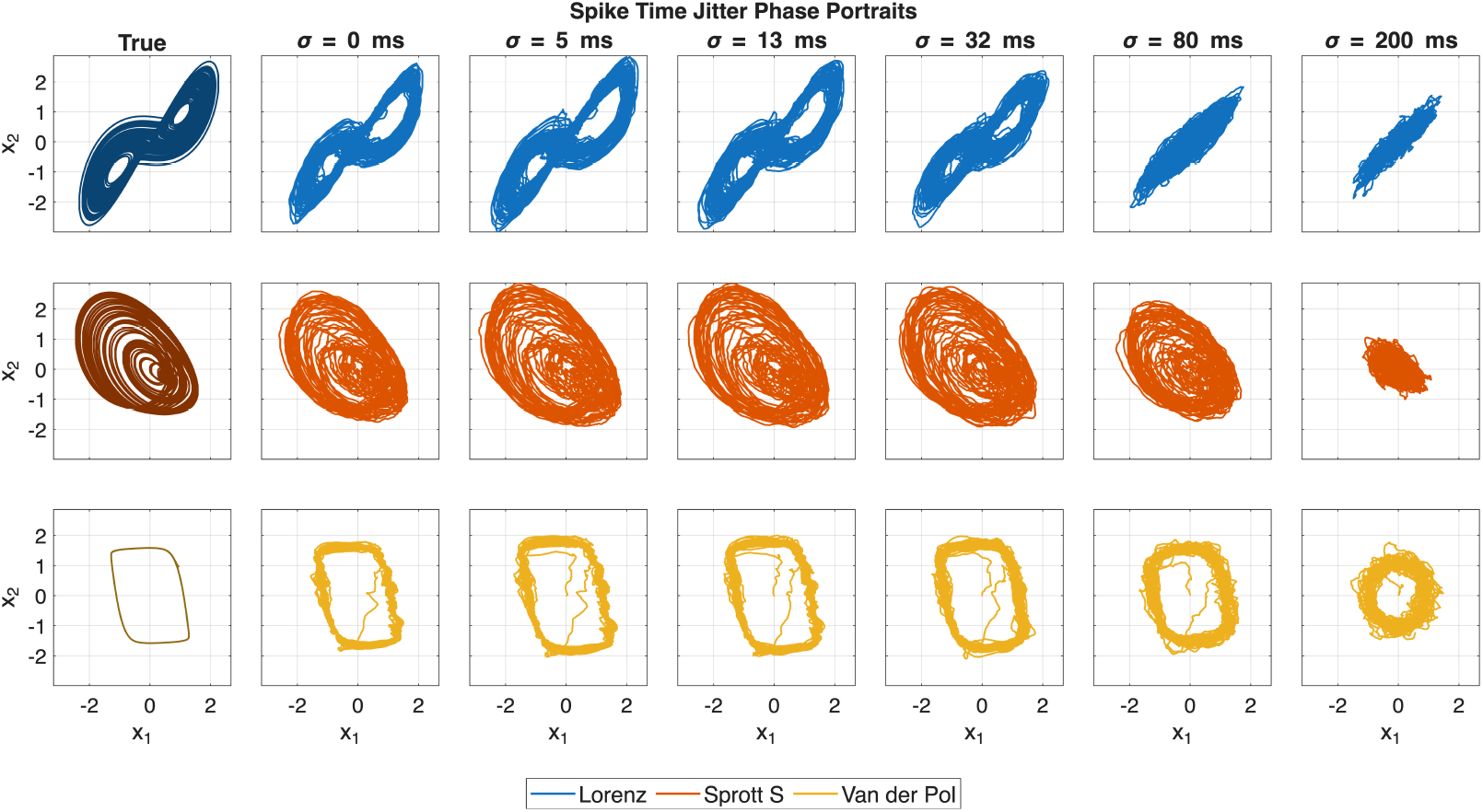
A spike-time jitter analysis of Trained BANFF Networks. The phase portraits for all dynamical systems for logarithmically spaced spike time jitters (*σ*) from 5 ms to 200 ms. All spike times were jittered randomly after closed-loop simulation of the dynamical system; the jitter magnitude for each spike was drawn from a Gaussian distribution with zero mean and a standard deviation of *σ*.

## Discussion

Neural circuits, both *in vivo* and *in vitro*, display considerable heterogeneity, particularly in the excitability of single neurons. Our central finding is that bias-only plasticity acting on the excitability of spiking neurons embedded in a single fixed recurrent network is sufficient to learn a broad repertoire of tasks in heterogeneous spiking networks. In continuous-time dynamical systems identification, the network reproduced key features (steady state attractors) of the target systems. In conventional supervised settings, the same SNN without any changes to recurrent synapses achieved competitive performance on a diverse set of classification and regression tasks. Together, these results show that adaptive control of excitability can tune network trajectories and output structure while recurrent connectivity remains static and each neuron is constrained to be exclusively either excitatory or inhibitory. Our results support recent work and perspectives on the importance of neural heterogeneity for computations.

This work extends our prior work, which considered rate-based neurons with static low-rank feedforward connectivity [19], to a recurrent spiking neural network with adaptive leaky integrate and fire neurons, along with the prior work of others in computing with artificial neural networks using bias modulation [17, 33]. We show that bias-only modulation of neuronal excitability continues to select and stabilise target vector fields under spiking dynamics. Together, these design choices provide a direct spiking instantiation of bias-only learning that supports rapid reconfiguration by switching learned bias sets, without retraining synapses or altering connectivity. Overall, this provides a more biologically plausible model of bias-learning than a rate-based network implementation.

Secondly, the inclusion of an adaptation current in the ALIF model endowed each neuron with a non-stationary, history-dependent input–output (*F*(*I*)) relationship, in which the instantaneous mapping from synaptic drive to spiking was dynamically gated by a slow variable *w* that integrated recent spiking activity. This slow feedback introduced an internal state that modulated excitability over time, such that each neuron’s responsiveness to a given input depended on its recent firing history [34, 35]. As a result, a learned bias current in such neurons was not a fixed offset, as in conventional rate-based networks, but acted as a control input whose effect was contingent on the current adaptation state. This represents a departure from rate-based formulations, in which a neuron’s transfer function is static and memoryless. The adaptive mechanism also opens the possibility for information to be encoded not only in average firing rates but also in the timing of spikes, since the neuron’s instantaneous excitability reflects a dynamic balance between input drive and adaptation [36]. While this does not guarantee the use of temporally precise codes, and we have shown that in this work a rate-based code persisted, it aligns more closely with biological neurons, which exhibit experimentally measured non-stationary *F*(*I*) curves and spike-frequency adaptation across multiple cell types [34, 37]. The ability of the bias-learning to achieve rapid learning and stable performance over several task types despite the dynamical complexity imposed by ALIF suggests that the BANFF framework has utility for controlling networks with challenging dynamical features.

Finally, we adopted a more biologically plausible learning rule than in our earlier BANFF work (which used backpropagation) by employing local, three-factor updates with eligibility traces in the spiking network. Each neuron maintains an eligibility trace that depends on its membrane variables and adaptation state, whilst a task-level broadcast error provides the modulatory factor; the product yields a bias update that is local in space and online in time, avoiding backpropagation-through-time [20]. This rule is compatible with neuromodulatory interpretations (e.g. dopamine-like signals) and supports learning over behavioural timescales while retaining BANFF’s key constraint of fixed synapses. In combination with the Δ*t*-invariant within-step alignment described above, the resulting scheme couples realistic single-neuron dynamics to a biologically grounded credit-assignment mechanism that is sufficient to configure closed-loop dynamics through bias modulation alone.

In contrast to our previous rate-based BANFF implementation, which exposed only a single linear decoder trained jointly with neuron-specific bias currents [19], the spiking BANFF network in this work introduces a secondary linear decoder that is driven by its own fixed synaptic cascade and trained without feeding back into the recurrent network core. This architecture can be interpreted as a minimal model of a downstream neural structure receiving projections from a cortical population: the recurrent spiking network learns task-appropriate trajectories through local three-factor updates acting on intrinsic excitability (biases), whilst the secondary decoder corresponds to efferent pathways that read out those trajectories with a higher signal-to-noise ratio or alternative temporal filtering. The separation between a fixed or slowly changing recurrent scaffold and flexible linear readouts is similar to reservoir-computing views of cortical microcircuits, in which random or low-rank recurrent dynamics are reused across many tasks and different linear combinations of the same population activity implement context-specific computations [15, 38]. Experimental work further supports the notion that behaviourally relevant variables can be recovered with high fidelity by downstream linear decoders, even when local covariance within the upstream population is approximately preserved during learning or when the underlying code drifts over days [38, 39]. Within this framework, our secondary decoder provides a biologically plausible surrogate for plastic readout pathways that can be trained or re-trained without perturbing the BANFF core, thereby allowing more accurate and task-specialised readout while retaining the key constraint that recurrent synaptic connectivity remains fixed.

BANFF was designed as a simplified model of cortical learning without synaptic plasticity to investigate the capabilities of a recurrent network with fixed random synaptic connections and endowed with plastic excitability. Synapses in the cerebral cortex, however, are not fixed. An overwhelming corpus of data indicates widespread synaptic plasticity in the cortex. To further develop this framework, learning of gated inputs/intrinsic excitability and within-cortex synaptic plasticity (e.g. through spike-timing-dependent plasticity) should be integrated into a cohesive model. Future work will focus on the integration of this effect into the BANFF, as well as the development of more biologically plausible models of the cortex and subcortical structures, such as the basal ganglia, which modulate information presented to cortical circuits [40, 41]. Such models should include a more realistic representation of different cell types (e.g. a correct E-I balance) as well as experimentally observed recurrent connection probabilities and modular networks representing cortical columns and different brain regions.

## Acknowledgments

WN is funded by an NSERC Discovery Grant, a Canada Research Chair, a Hotchkiss Brain Institute start-up grant and the Cumming Medical Research Fund. AG is supported by NSERC, Digital Research Alliance of Canada, Hotchkiss Brain Institute, Alberta Children’s Hospital Research Institute, and the Azrieli Accelerator. This research was partially funded by Synaptrain Technologies Inc.

## Author Contributions Statement

WN and AG supervised the project. KM and NA performed numerical simulations. KM performed mathematical analysis. KM, WN and AG prepared and edited the manuscript.

## Code availability

Code for this work will be made available at the following location: https://github.com/kaimason100/BANFF_SNN/tree/main

## Methods

Numerical values of all simulation parameters are listed in Table 2.

**Table 2:** Common SNN hyperparameters across all tasks. Numerical parameters pertaining to the SNN that were shared by all simulations.

| Parameter | Value | Notes |
| --- | --- | --- |
| **Discretisation & event alignment** |  |  |
| Time step $\Delta t$ | 1 ms | Used for all tasks. |
| **ALIF neuron model** |  |  |
| Membrane time constant $\tau_u$ | 50 ms | $\alpha = e^{-\Delta t / \tau_u}$. |
| Adaptation time constant $\tau_w$ | 500 ms | $\beta = e^{-\Delta t / \tau_w}$. |
| Leak potential $E_L$ | -70 mV | |
| Spike threshold $V_{th}$ | -50 mV | |
| Reset potential $V_{reset}$ | -65 mV | |
| Spike-triggered increment $b$ | 0.5 mV | |
| **Surrogate gradient** |  |  |
| Surrogate $\psi(\tilde{u})$ form | triangular | Evaluated at LSTI crossing (pre-reset $\tilde{u}$). |
| Surrogate slope $\phi_u$ | 1 | |
| Surrogate half-width $\delta_u$ | 0.8 mV | |
| **Synaptic cascades** |  |  |
| Primary rise constant $\tau_{s, rise}$ | 2 ms | Two-stage $(x, r)$; $r_k$ feeds primary head and recurrent block. |
| Primary decay constant $\tau_{s, decay}$ | 50 ms | " " |
| Secondary decoder rise constant $\tau_{rise}^{aux}$ | 10 ms | Drives secondary decoder; no feedback into core. |
| Secondary decoder decay constant $\tau_{decay}^{aux}$ | 200 ms | " " |
| **Architecture & scaling** |  |  |
| Low-rank recurrent matrix dimension | 10 | Base decoder width; $R = \max$ task output dimension. |
| Excitatory fraction $p_{exc}$ | 0.5 | Dale's law via column-wise rectification; $\eta \geq 0$. |
| Recurrent gain $S_{rec}$ | 0.05 | Scales $\eta h_k$; instantaneous self-term compensated. |
| Primary decoder gain $S_{dec}$ | 0.1 | $W_{out} = S_{dec} \cdot W_{out}^{base}[1:D_{out}, :]$. |
| Encoder gain $S_{enc}$ | 2 | Magnitude scale of $W_{in}$ entries. |
| Input scaling | $1/\sqrt{D_{in}}$ | Applied after $z$-scoring (size-invariant drive). |

### Spiking neuron model

We used a discrete–time ALIF model with a spike–triggered, hyperpolarising adaptation current and within–step LSTI.

For hidden neuron *j* at time step *k*, the pre–reset voltage obeys

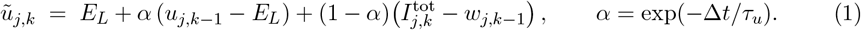

A spike is emitted if *ũ*_*j,k*_ ≥ *V*_th_, i.e.

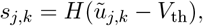

followed by a hard reset

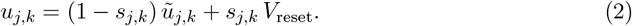

Spike–frequency adaptation is purely spike–triggered:

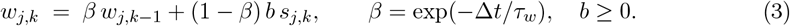

We used the two–stage synaptic cascade described below, implemented in an LSTI–consistent form that is approximately Δ*t*–invariant. In the absence of a spike within the step (*s*_*j,k*_ = 0), the single–segment decays reduce to

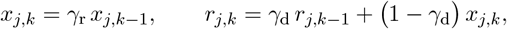

with *γ*_r_ = exp(− Δ*t*/*τ*_*s*,rise_) and *γ*_d_ = exp(− Δ*t*/*τ*_*s*,decay_), where *τ*_*s*,rise_ and *τ*_*s*,decay_ are the synaptic rise and decay time constants, respectively. Here *x*_*j,k*_ encodes the rise and *r*_*j,k*_ the effective synaptic drive.

If a spike occurred within step *k* at fraction *ρ* ∈ [0, 1] (linear interpolation of *u* between *k* − 1 and *k*), we split the step into a pre–segment of duration *ρ*Δ*t* and a post–segment (1 − *ρ*)Δ*t*. Let 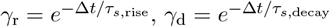 and define fractional decays

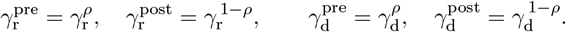

With no continuous drive between events, the pre–segment advances as

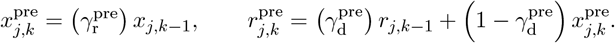

At the spike time, the rise state receives a fixed, dt-invariant impulse

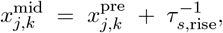

followed by the post–segment updates

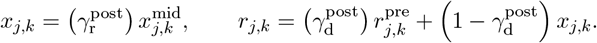

If there was no spike (*s*_*j,k*_ = 0), we used the single–segment decays above. All other states (*u, w*) used the same pre/post split, and the surrogate gradient *ψ*(*ũ*_*j,k*_) was evaluated at the pre–reset crossing potential *ũ*_*j,k*_.

### Network architecture

We adopted a fixed–synapse, low–rank recurrent architecture for the spiking core together with linear decoders. Let *D*_in_ be the input feature dimension, *D*_out_ the output dimension of the primary head for the current task, and *N* the number of hidden neurons. At step *k*, the hidden activity is the synaptic trace vector *r*_*k*_ ∈ ℝ^*N*^. Inputs were scaled by 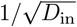 for size invariance and embedded by

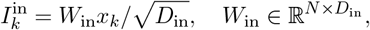

where 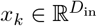 denotes the input vector at time step *k* (Table 3).

**Table 3:** Dataset statistics. The respective sizes of the training, validation and test datasets for each task, as well as the expected (exp.) and actual (act.) numbers of training epochs. The expected number of training epochs was used to calculate the rate of decay for the bias learning rate (Equation 13).

| Task/system | $N_{\text{train}}$ | $N_{\text{val}}$ | $N_{\text{test}}$ | Epochs (exp.) | Epochs (act.) | Notes |
| --- | --- | --- | --- | --- | --- | --- |
| **Dynamical systems** |  |  |  |  |  |  |
| Lorenz | – | – | – | 200,000 | 200,000 | 10 s supervisor windows sampled from a 2000 s trajectory (new window each epoch). Alternating open- (30 ms) and closed-loop (55 ms, off-policy) training. System simulated at $2 \times$ speed. Closed-loop testing. |
| Sprott S | – | – | – | 200,000 | 200,000 | 10 s supervisor windows sampled from a 2000 s trajectory (new window each epoch). Alternating open- (30 ms) and closed-loop (55 ms, off-policy) training. System simulated at $4 \times$ speed. Closed-loop testing. |
| Van der Pol | – | – | – | 200,000 | 200,000 | 5 s supervisor windows sampled from a 2000 s trajectory (new window each epoch). Alternating open- (30 ms) and closed-loop (55 ms, off-policy) training. System simulated at $8 \times$ speed. Closed-loop testing. |
| **Classification** |  |  |  |  |  |  |
| Breast cancer | 341 | 113 | 115 | 500 | 500 | Each sample presented for 300 ms (300 time steps). |
| MNIST | 6,000 | 2,000 | 2,000 | 500 | 2,000 | Each sample presented for 300 ms (300 time steps). Training extended beyond the expected number of epochs to ensure convergence. |
| Fashion-MNIST | 42,000 | 14,000 | 14,000 | 500 | 209 | Each sample presented for 300 ms (300 time steps). Training stopped early due to early convergence. |
| **Regression** |  |  |  |  |  |  |
| Car price | 4,042 | 1,347 | 1,349 | 500 | 800 | Each sample presented for 300 ms (300 time steps). Training extended beyond the expected number of epochs to ensure convergence. |
| Abalone age | 2,506 | 835 | 836 | 500 | 1000 | " " |
| Yacht hydrodynamics | 197 | 49 | 62 | 500 | 500 | Each sample presented for 300 ms (300 time steps). |

We set the total somatic drive as

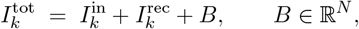

with the bias *B* trained while all synaptic matrices remained fixed. Component-wise we write 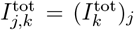. For each task, *D*_out_ was chosen to match the required output dimension: the state dimension of the dynamical system, the number of classes for classification problems, or *D*_out_ = 1 for scalar regression.

Recurrent drive used a low–rank factorisation. We defined a base decoder 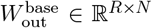 with small rank *R* = 10, chosen such that *R* = max_tasks_(*D*_out_), and a non–negative matrix *η* ∈ ℝ^*N*×*R*^. Projected activity

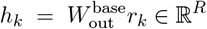

was used to produce the recurrent drive

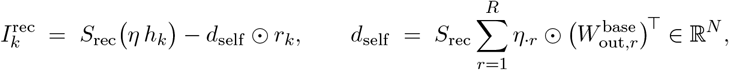

with scalar gain *S*_rec_. The vector *d*_self_ is precisely the diagonal of the effective low–rank recurrent matrix 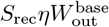 its *j*th entry is 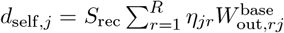. Subtracting *d*_self_ ⊙ *r*_*k*_ therefore cancels the instantaneous self–coupling introduced by the shared encoder/decoder structure.

To impose Dale’s law, for each presynaptic neuron *j*, we sampled a sign *σ*_*j*_ ∈ {+1, −1} with ℙ [*σ*_*j*_ = +1] = *p*_exc_ = 0.5. We then rectified the columns of 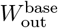 by setting

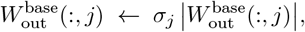

and constrained *η* ≥ 0. With this construction, all outgoing synapses from the presynaptic neuron *j* share the same sign *σ*_*j*_ (Dale’s law), while the low–rank structure and self–coupling compensation are preserved.

A fixed decoder selected *D*_out_ rows of 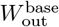 and applied a scalar scale,

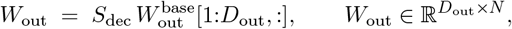

to produce the primary head *z*_*k*_ = *W*_out_*r*_*k*_. For evaluation/selection, we also used a downstream secondary readout 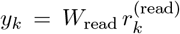 driven by its own two–stage cascade 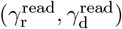 with fixed time constants (Table 2). This secondary head never fed back into the recurrent dynamics.

### Learning algorithm

We used e–prop [20] with bias–only plasticity in the recurrent core: the input weights *W*_in_, the fixed recurrent readout block 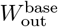, the matrix *η*, and the primary decoder *W*_out_ were fixed; only neuron biases *B* ∈ ℝ^*N*^ were trained. A secondary readout *W*_read_ was optimised for evaluation and did not influence the recurrent dynamics.

At discrete time *k* ∈ {1, …, *T* }, the network produces the primary output

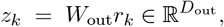

where *r*_*k*_ ∈ ℝ^*N*^ denotes the synaptic decay–stage state (the second stage of a two–stage rise–decay synaptic cascade). Given a target 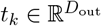 and a per–step loss *ℓ*_*k*_ = *ℓ*(*z*_*k*_, *t*_*k*_) (e.g. mean–squared error or cross–entropy), the decoder–space gradient is

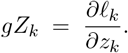

Each neuron *j* then received a broadcast learning signal

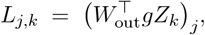

and maintained a local bias–eligibility 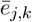 (defined below). The instantaneous contribution to the bias gradient was the three–factor product

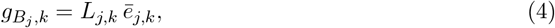

and the online e–prop estimate of the sequence gradient was

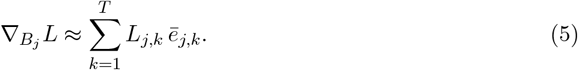

We computed *L*_*j,k*_ from the readout gradient at discrete time–step *k*, and aligned the neuron–local eligibility 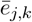 to the same within–step crossing state as the surrogate (see below), using the pre–reset potential *ũ*_*j,k*_ and the LSTI split. Because spikes were non-differentiable, the spike derivative was replaced by a piecewise–linear surrogate evaluated at the inferred within–step crossing. Let Δ*t* denote the simulation step, *V*_th_ the threshold, and *ũ*_*j,k*_ the pre–reset membrane potential at the crossing time obtained by LSTI. The surrogate factor was

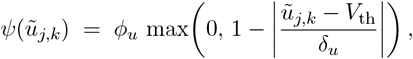

with slope *ϕ*_*u*_ and window half–width *δ*_*u*_. This factor gated the raw neuron–local eligibility and thus entered the learning rule via

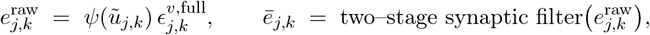

where 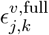 denotes the membrane sensitivity to the bias at the pre–reset crossing (including spike–triggered adaptation). The filtered term 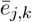 is the eligibility used in 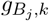 above.

LSTI returned a fractional spike time *ρ*_*j,k*_ ∈ [0, 1] within the step. To ensure temporal alignment and Δ*t*–invariance, all exponentials were split at this fraction. For the membrane and adaptation decays, we defined

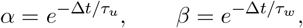

and used

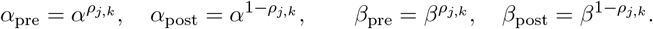

Thus, 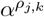 represents the membrane decay over a subinterval of duration *ρ*_*j,k*_Δ*t*, and 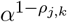 represents the decay over the remaining subinterval (1 − *ρ*_*j,k*_)Δ*t*. For the synaptic cascade, we similarly defined

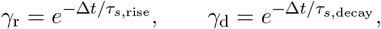

and applied the same pre/post split to (*γ*_r_, *γ*_d_). Full pre/reset/post eligibility recursions and synaptic–filter updates (sufficient for reimplementation) are provided in the Supplementary Material.

Collecting the terms above, the bias update can be written compactly in vector form. Let 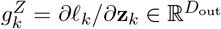 denote the decoder-space gradient at time step *k*, and define the broadcast learning signal and eligibility vectors

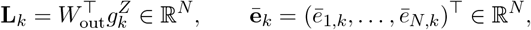

where *N* is the number of neurons in the recurrent core. For a single sequence of length *T* with total loss 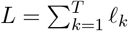, the e-prop estimator of the gradient of *L* with respect to the bias vector **B** ∈ ℝ^*N*^ is

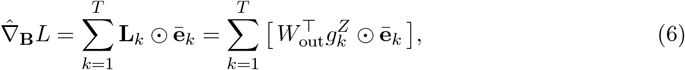

where ⊙ denotes the element-wise (Hadamard) product. For a mini-batch of *S* sequences, we average these contributions to obtain

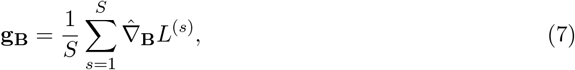

and use **g**_**B**_ as the gradient input to the Adam optimiser described in the Optimisation subsection, yielding a parameter update of the form

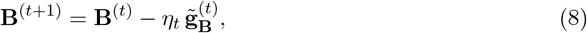

where 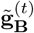 is the Adam-preconditioned version of **g**_**B**_ at iteration *t* and *η*_*t*_ is the learning rate. Thus, each bias component is updated by a three-factor rule combining a task-level learning signal, a neuron-local eligibility trace, and the optimiser’s adaptive preconditioning.

### Dynamical Systems Tasks

We considered general *D*–dimensional systems 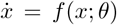 and trained the network for one-step prediction. Reference trajectories were generated numerically by Euler integration. We computed global per–dimension mean and standard deviation (*µ*_global_, *σ*_global_) on a long reference run and applied fixed *z*–scaling to all splits:

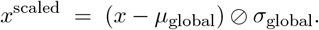

During training, we sampled contiguous windows of duration *T*_sim_ (with *K* = ⌊*T*_sim_*/*Δ*t*⌋ steps) uniformly from a precomputed simulation of 2000 s, drawing a fresh window each epoch. The loss was defined as the one–step mean–squared error on the fixed primary head,

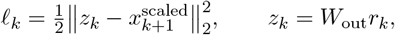

and the sequence loss was *L* = ∑_*k*_ *ℓ*_*k*_.

Training used an alternating supervision schedule repeated across the window:

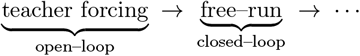

In teacher–forcing steps, inputs were the truth 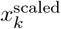; in free–run steps, inputs were the model’s own *z*_*k*−1_, while the supervisor remained the ground–truth 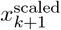 (off–policy). In all reported results, we used fixed segment lengths of 30 ms teacher forcing followed by 55 ms free–run (with Δ*t* = 1 ms) (Figure 1E).

All bias–parameter gradients were accumulated online via the e–prop three–factor rule.

Every 50 epochs, we paused training to run diagnostic closed-loop rollouts (no gradient flow). From five initial conditions, we generated 50 s rollouts under both heads and computed geometry-aware statistics as detailed below.

For each dynamical system, we selected up to three 2-D state pairs (*x*_*a*_, *x*_*b*_) (for example, (*x*_1_, *x*_2_), 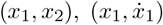, or 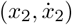). For each chosen pair, we formed point clouds from the model (*P*) and the ground truth (*Q*) by time–subsampling every *s*_sub_ = 5 simulation steps, to reduce redundancy between consecutive states. We then quantified the discrepancy between *P* and *Q* using a trimmed sliced 2–Wasserstein distance. For each random direction, we define

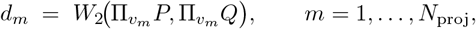

and apply the trimming to the collection of per–direction distances:

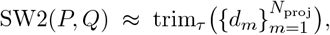

where 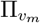 denotes projection onto a random unit direction *v*_*m*_ ∈ 𝕊^1^, *W*_2_ is the one-dimensional 2-Wasserstein distance between the projected sets (computed by matching empirical quantiles of the sorted projected points), *N*_proj_ = 256, and trim_*τ*_ is a symmetric trimmed mean with fraction *τ* = 0.10 (we discard the largest and smallest 10% of per–direction distances before averaging).

In more concrete terms, for each random direction *v*_*m*_ we take a ‘shadow’ of both point clouds along that line (by projection), sort the resulting one-dimensional samples, compute how far the two sorted lists differ on average (the 1-D Wasserstein distance), and then average these discrepancies over many directions. The trimming step makes this average robust by removing directions that produce unusually small or large discrepancies (for example, nearly degenerate alignments). We finally report the SW2 value averaged over the selected 2-D state pairs. See the Supplementary Material for full details of the SW2 calculation.

At the end of training, for each head separately, we selected the epoch index that minimised the phase portrait Wasserstein distance over the diagnostic snapshots taken every 50 epochs. The corresponding biases and secondary readout weights were retained for final reporting.

Using the selected biases and readout weights, we simulated the trained network in an autonomous closed loop. For each dynamical system, we first drew a single random initial condition. Starting from this state, we ran a 20 s warm-up of the closed-loop network and, in parallel, a 20 s simulation of the corresponding continuous-time dynamical system at the same time step Δ*t* (and with the appropriate rate parameter). At the end of the warm-up, the network state was used as the initial condition for a fresh 50 s simulation of the ground-truth system, while the network was simply continued for a further 50 s without resetting. We reported (i) phase portraits, (ii) Poincaré sections, and (iii) return maps computed from these main 50 s segments only.

For each dynamical system, we used the closed-loop trajectories of both the trained network (secondary readout) and the corresponding ground-truth system, stored as time series *x*_read_(*t*) and *x*_true_(*t*) with *D* state dimensions. All visualisations were computed from these precomputed trajectories using a time window of 50 s.

Let *x*(*t*) ∈ ℝ^*D*^ denote either *x*_read_(*t*) or *x*_true_(*t*) evaluated at the indices ℐ. For each system, we formed the set of coordinate pairs

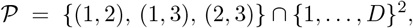

i.e. (*x*_1_, *x*_2_) for the Van der Pol oscillator (*D* = 2) and (*x*_1_, *x*_2_), (*x*_1_, *x*_3_), (*x*_2_, *x*_3_) for the three– dimensional systems (Lorenz and Sprott S). For each (*a, b*) ∈ 𝒫 we plotted the network trajectory (*x*_read,*a*_(*t*), *x*_read,*b*_(*t*)) and the ground–truth trajectory (*x*_true,*a*_(*t*), *x*_true,*b*_(*t*)) in separate, side–by– side panels.

Poincaré sections were computed only for the three-dimensional systems (Lorenz and Sprott S), for which we defined one or more axis-aligned sections of the form

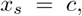

where *s* ∈ {1, …, *D*} is the section coordinate, *c* is the section level, and a sign constraint on the crossing velocity is specified by dir ∈ [+1, − 1}. For each section and each trajectory (network and ground truth separately), we considered consecutive samples *x*(*t*_*k*_), *x*(*t*_*k*+1_) in the chosen time window and detected candidate crossings whenever

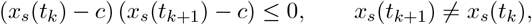

and the sign of the increment *x*_*s*_(*t*_*k*+1_) − *x*_*s*_(*t*_*k*_) matched the prescribed direction `dir`.

For each accepted crossing we linearly interpolated between *t*_*k*_ and *t*_*k*+1_ to estimate the intersection time

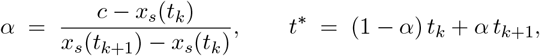

clipped to *α* ∈ [0, 1], and computed the corresponding state *x*(*t*^∗^) by linear interpolation in every coordinate. For each section coordinate *s*, we then projected these intersection points onto all remaining coordinate pairs (*a, b*) ∈ 𝒫 with *a*≠ *s, b*≠ *s*, yielding Poincaré point clouds in (*x*_*a*_, *x*_*b*_).

Classical peak return maps were constructed for the three-dimensional systems (Lorenz and Sprott S). For each system and each state coordinate *d* ∈ {1, …, *D*}, we extracted the corresponding scalar time series *x*_*d*_(*t*) over the selected time window for both the network and the ground–truth trajectories. We then identified local maxima using a standard peak–finding routine: MATLAB’s `findpeaks` was applied with a minimum peak prominence of 1. This yielded sequences of peak values 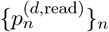and 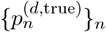 for the network and ground truth, respectively.

For each coordinate *d*, we formed the one–step return pairs

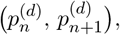

for all *n* such that both peaks exist, and plotted these as scatter points with 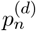 on the horizontal axis and 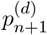 on the vertical axis.

### Classification tasks

For a *C*-class problem, we created random train/validation/test splits (60/20/20). Features 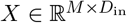 were *z*–scored with training–set statistics,

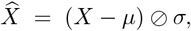

and labels were mapped to one–hot vectors *y* ∈ {0, 1}^*C*^ (per sample). Inputs to the spiking core were further scaled by 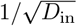 to make the effective drive approximately size-invariant (Table 3).

Each sample was presented for 300 ms with Δ*t* = 1 ms. Let *K* = ⌊0.300/Δ*t*⌋, *K*_avg_ = ⌊0.5 *K*⌋, and *k*_0_ = *K* − *K*_avg_ + 1. Per-step logits for the two heads were

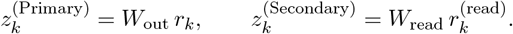

We averaged the last 50% of the window:

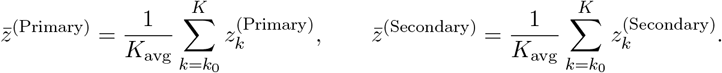

The primary–head loss was cross–entropy with 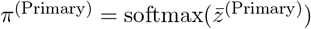:

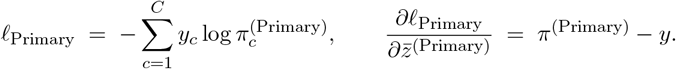

The secondary readout was trained with the same target, without feedback into the core:

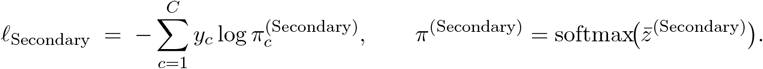

Per–sample e–prop gradients (from *ℓ*_Primary_ via the fixed decoder *W*_out_) were accumulated within an epoch to update *B* once per epoch by Adam with AMSGrad and an exponentially decaying learning rate; *ℓ*_Secondary_ updated *W*_read_ only. Model selection used the highest validation accuracy. Test metrics were computed once using the selected checkpoint.

### Regression tasks

Given samples(*x*^(*i*)^,*y* ^(*i*)^)with 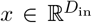 and *y* ∈ ℝ, we created random train/validation/test splits (60/20/20). Inputs were *z*–scored with training–set statistics and scaled by 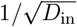 before the encoder (Table 3). Targets were *z*–scored (using training–set mean and s.d.) for optimisation; when reporting in original units, we inverted this transform.

Each sample was presented for 300 ms with Δ*t* = 1 ms. Using the same indices *K, K*_avg_, *k*_0_ as above, per–step logits and their averages were

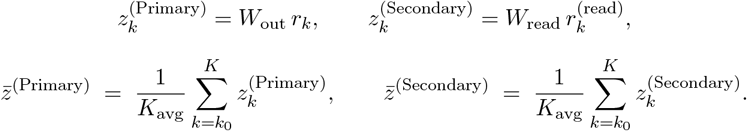

Mean-squared errors for the two heads were

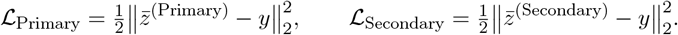

Per–sample e–prop gradients (from ℒ _Primary_ via *W*_out_) were accumulated within an epoch to update *B* once per epoch by Adam with AMSGrad and an exponentially decaying learning rate; ℒ _Secondary_ updated *W*_read_ only. Alongside loss, we tracked Pearson correlation *r* between predictions and targets on each split. Model selection used the highest validation correlation.

### Optimisation

Network parameters were trained using Adam [42] with the AMSGrad [43] variant to ensure non-increasing second-moment estimates. For each parameter vector *θ* (biases *B* for the spiking neurons and the linear readout weights *W*_read_ for the secondary head), we computed gradients *g*_*t*_ at iteration *t* and updated the biased first- and second-moment estimators as

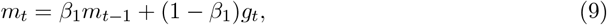

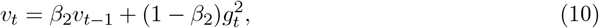

with *β*_1_ = 0.9, *β*_2_ = 0.999 and *ε* = 10^−8^. AMSGrad replaces the standard Adam second-moment term with a monotonically non-decreasing estimator,

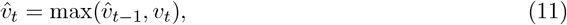

and parameters are updated according to

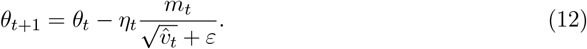

Bias parameters *B* used a decaying learning rate *η*_*t*_ = *η*(*e*) determined at the epoch level. We employed an exponential schedule defined by

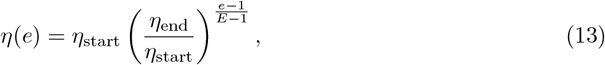

where *e* is the epoch index (1-indexed). Here *η*_start_ = 5 × 10^−1^ and *η*_end_ = 5 × 10^−3^ for classification and regression tasks and *η*_start_ = 5 × 10^−2^ and *η*_end_ = 5 × 10^−3^ for dynamical systems tasks. The parameter *E* denotes the expected total number of training epochs for the task and varies between tasks. This value differed between tasks because the anticipated training horizon varied across experiments; the learning-rate decay was therefore completed at different epoch numbers depending on the task (Table 3). For epochs beyond *E*, the same functional form was applied, resulting in learning rates lower than *η*_end_. The total number of executed training epochs was chosen heuristically based on observation of the training performance.

The secondary head parameters *W*_read_ used a fixed learning rate of 1 × 10^−4^ throughout training.

### Spike-time Jitter Analysis

To assess whether the readout representation of the trained BANFF spiking network implemented predominantly rate- or time-based codes for the dynamical-systems tasks, we performed a post hoc spike-time jitter analysis on the closed-loop trajectories. For each system (Lorenz, Sprott S and Van der Pol), we first ran a closed-loop simulation of the trained network using the task-specific bias set and secondary readout weights. The network was simulated with a fixed time step Δ*t* = 1 ms, and all analyses were performed on the secondary head driven by its own two-stage synaptic cascade (rise/decay time constants 10 ms and 200 ms, respectively).

During the closed-loop rollout we recorded, for every hidden neuron, the set of discrete time-step indices *k* ∈ {1, …, *K* − 1} at which at least one spike occurred, where *K* is the number of samples in the analysis window. Each such index was treated as the time of a spike event occurring within the corresponding interval [*k*Δ*t*, (*k* + 1)Δ*t*). Let *k*_*i,n*_ denote the *n*-th spike index of neuron *i*.

We then generated jittered spike trains by independently perturbing each spike index by a Gaussian-distributed time offset. Specifically, for a given jitter standard deviation *σ* (in seconds), we drew an integer-valued offset

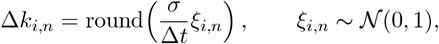

and formed the jittered index

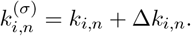

Indices that moved outside the analysis window were discarded, i.e. we enforced

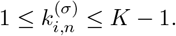

To avoid introducing multiple spikes within a single time step for a given neuron, any remaining duplicates were merged by retaining only the first occurrence of each index. In this way, spike times were perturbed at the granularity of the numerical integration step, with an effective temporal jitter whose standard deviation is, up to discretisation effects, equal to *σ*. We evaluated *σ* ∈ {0, 5, 13, 32, 80, 200} ms, corresponding to a logarithmically spaced set between 5 ms and 200 ms plus a zero-jitter baseline (*σ* = 0 leaves the spike trains unchanged).

For each jitter level, we reconstructed a secondary-head trajectory by replaying the jittered spike trains through the fixed double-exponential synaptic filter and secondary readout matrix. The recurrent core was not re-simulated; only the readout pathway was recomputed from the modified spikes, ensuring that any changes in the output arose purely from spike-time perturbations rather than altered recurrent dynamics. From the resulting time series, we computed phase portraits in the same z-scored coordinates and over the same 50 s windows used for the unjittered closed-loop analysis.

### Computational implementation

All simulations were implemented using custom code in MATLAB on a desktop and/or workstation computer. Some simulations utilised CUDA code (callable from MATLAB with MEX) for a GPU-driven speed-up.

## Supplementary Material

### Eligibility trace for an ALIF neuron

We derive here the e–prop eligibility trace for bias–only learning in an adaptive leaky integrate–and–fire (ALIF) neuron with purely spike–triggered adaptation. All updates are aligned to the inferred within–step spike time using linear spike–time interpolation (LSTI).

Let the simulation time–step be Δ*t*, and define the discrete decay factors

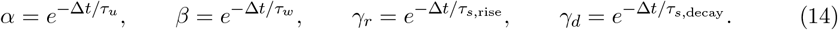

For a single hidden neuron (index *j* suppressed), with membrane potential *u*_*k*_, adaptation current *w*_*k*_, and spike *s*_*k*_ ∈ {0, 1} at discrete time *k*, the forward dynamics are

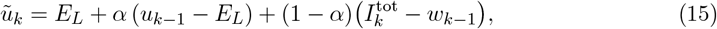

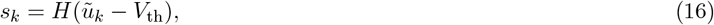

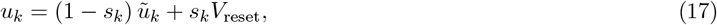

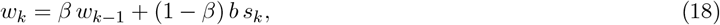

where *E*_*L*_ is the leak reversal potential, *V*_th_ the threshold, *V*_reset_ the reset potential, and *b >* 0 is the strength of spike–triggered adaptation. Spikes are filtered by a two–stage synaptic cascade,

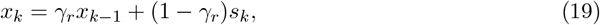

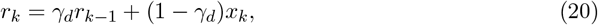

and the readout is *z*_*k*_ = *W*_out_*r*_*k*_ with fixed *W*_out_.

With per–step loss *ℓ*_*k*_ = *ℓ*(*z*_*k*_, *t*_*k*_) and output–space gradient 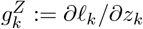, the neuron–specific broadcast learning signal is

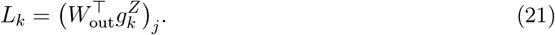

We consider bias–only learning in the recurrent core, with an additive bias parameter *B* entering the total input as 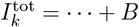, so that 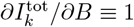.

Let 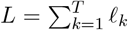 be the total loss over a sequence. The e–prop identity factorises the gradient of *L* with respect to *B* into a product of a learning signal and a neuron–local eligibility trace,

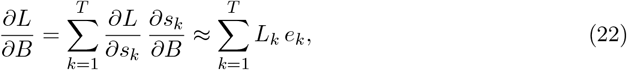

where *e*_*k*_ is the eligibility associated with the spike *s*_*k*_, and the broadcast signal *L*_*k*_ approximates ∂*L/*∂*s*_*k*_.

The hard threshold nonlinearity *H*(·) is replaced by a surrogate derivative evaluated at the pre–reset potential *ũ*_*k*_,

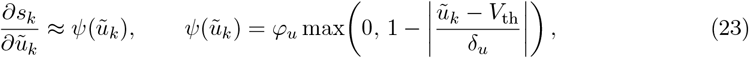

with slope *φ*_*u*_ and width *δ*_*u*_. By the chain rule, this gives

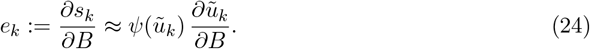

We therefore need ∂*ũ*_*k*_/∂*B* at the inferred crossing time inside the time–step. This is obtained from internal eligibility states that track the sensitivity of the membrane and adaptation variables to *B*.

Given (*u*_*k*−1_, *ũ*_*k*_), we infer a continuous threshold crossing within step *k* by

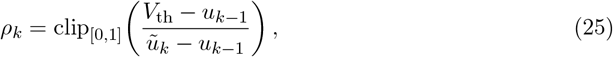

and split the leaky recursions into a pre–segment of duration *ρ*_*k*_Δ*t* and a post–segment of duration (1 − *ρ*_*k*_)Δ*t* using fractional decays

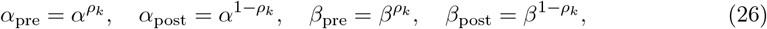

and analogously 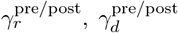 for the synaptic filter. Surrogate derivatives are evaluated at the pre–reset crossing potential *ũ*_*k*_, which is the LSTI interpolation point. When differentiating the LSTI–split recursions below, we treat *ρ*_*k*_ and the associated fractional decay factors as constants with respect to the bias parameter *B*; that is, we neglect derivatives of the inferred crossing time with respect to *B*, which is a standard approximation in surrogate–gradient treatments of spiking neurons.

We maintain two internal eligibility states at the end of step *k*:

- 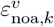 membrane eligibility that ignores feedback through adaptation, approximating ∂*u*_*k*_/∂*B* if *w* were clamped;
- 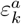 adaptation eligibility, approximating ∂*w*_*k*_/∂*B*.

Before the spike within step *k*, the membrane evolves (over the pre–segment) as

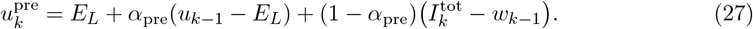

Differentiating with respect to *B* while temporarily treating *w*_*k*−1_ as constant gives the pre–segment membrane eligibility that ignores adaptation:

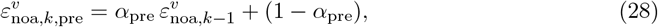

Because 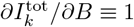. The adaptation current satisfies

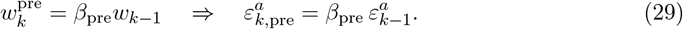

To obtain the full sensitivity of the pre–reset membrane potential, we must also account for the inhibitory contribution of *w*_*k*−1_ to *ũ*_*k*_ in (15). Differentiating the term −(1 − *α*_pre_)*w*_*k*−1_ yields

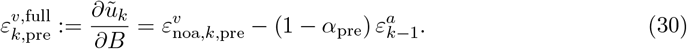

The raw spike eligibility at the inferred crossing time is therefore

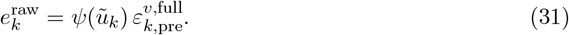

At the inferred spike time within step *k* (if *s*_*k*_ = 1), the membrane is reset to *V*_reset_ and the adaptation current receives a jump of size *b*:

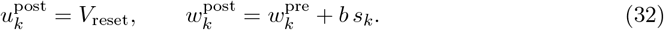

Since 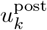 no longer depends on *B*, we obtain

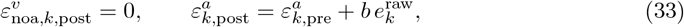

where the second relation follows from differentiating the adaptation update 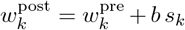 and using 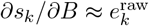.

From just after the spike to the end of the step, the membrane and adaptation again evolve leakily over the post–segment:

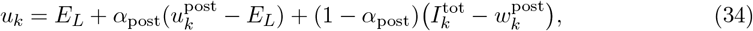

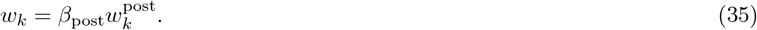

Differentiating with respect to *B* (and again using 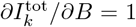) gives the end–of–step recusions

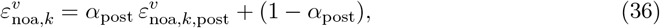

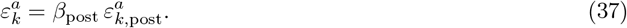

When *ρ*_*k*_ ∈ {0, 1} (no within–step crossing), the fractional decays collapse to either (*α*, 1) or (1, *α*) (and similarly for *β*), and the above expressions reduce to the usual single–segment recursions.

The postsynaptic variable *z*_*k*_ depends on past spikes only through the synaptic trace *r*_*k*_. Since *r*_*k*_ is obtained from *s*_*k*_ by a linear double–exponential filter, the derivative ∂*r*_*k*_/∂*B* obeys the same linear dynamics as *r*_*k*_ when driven by the raw eligibility 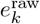 instead of *s*_*k*_. We therefore define rise– and decay–stage filtered eligibilities 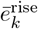 and 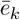, and update them using the same LSTI split as for *x*_*k*_ and *r*_*k*_.

For *s*_*k*_ = 1 (a spike within the step), we first propagate the previous values to the pre– segment end:

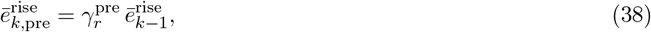

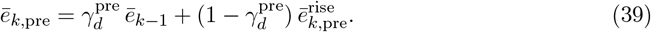

At the inferred spike time, we inject an impulse proportional to 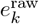 into the rise–stage eligibility,

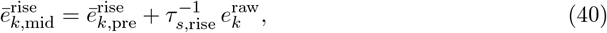

and propagate to the end of the step:

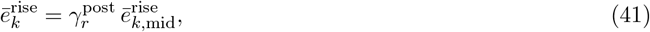

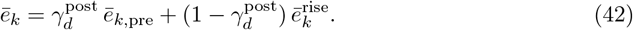

The fixed impulse 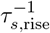 yields a jump size that is invariant under changes in Δ*t*. If *s*_*k*_ = 0 (no spike within the step), we use the standard single–segment updates

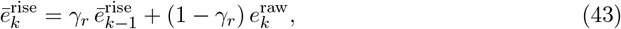

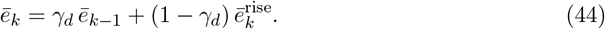

The filtered eligibility 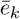 is the quantity that matches how perturbations of *B* influence the readout *z*_*k*_ through the synaptic cascade. The e–prop estimator of the bias gradient is then

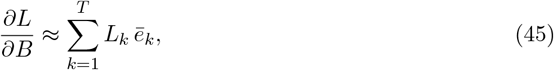

and, when the loss depends on a time–average 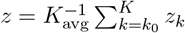 over a final window, the same filtered eligibility enters the window–averaged gradient as in the main text.

### Sliced Wasserstein distance (SW2)

We quantify discrepancies between planar point clouds (for example, phase portraits, Poincaré section points, or return–map pairs) using the sliced 2-Wasserstein distance. Let 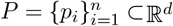 and 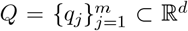 be finite point clouds (for our phase–portrait applications, *d* = 2).

We associate with these point clouds the empirical measures

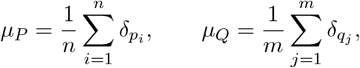

where *δ*_*x*_ is the Dirac measure at *x*.

For a unit vector *θ* ∈ *S*^*d*−1^ (the unit sphere in ℝ^*d*^), the push forward measure ⟨*θ, µ*⟩ is the distribution of the scalar projection *θ*^⊤^*x* when *x* ~ *µ*. The population sliced 2–Wasserstein distance between *µ*_*P*_ and *µ*_*Q*_ is then defined as

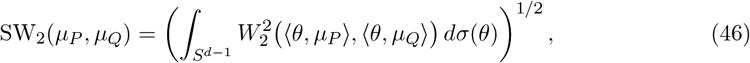

where *W*_2_ is the one–dimensional quadratic Wasserstein distance and *σ* is the uniform probability measure on the unit sphere *S*^*d*−1^.

In practice, we approximate the integral over directions by Monte Carlo sampling. We draw *K* i.i.d. random vectors *v*_*k*_ ~ 𝒩 (0, *I*_*d*_) and normalise them onto the unit sphere,

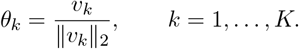

For a fixed direction *θ*, we project all points in *P* and *Q* onto this line and sort the resulting scalars:

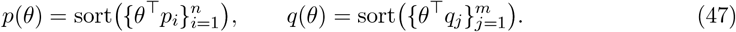

These sorted lists are empirical samples from the one-dimensional projected distributions ⟨*θ, µ*_*P*_⟩ and ⟨*θ, µ*_*Q*_⟩.

If the sample sizes differ (*n* ≠ *m*), we interpolate the empirical quantile functions onto a common grid. Let

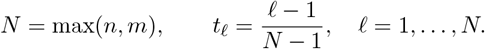

We form interpolated empirical quantile functions 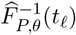 and 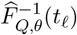 for the projected distributions using linear interpolation on the sorted samples *p*(*θ*) and *q*(*θ*). The slice–wise empirical 2-Wasserstein distance along direction *θ* is then approximated by

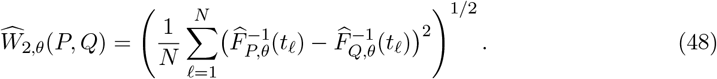

We compute these slice–wise distances for the *K* random directions,

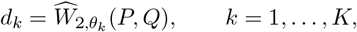

and sort them increasingly, *d*_(1)_ ≤ · · · ≤ *d*_(*K*)_. To reduce the influence of outlying directions, we use a trimmed mean. For a trimming fraction *τ* ∈ [0, 1/2), define *κ* = ⌊*τ K*⌋. The trimmed sliced 2-Wasserstein distance is then

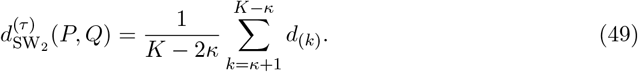

Unless otherwise stated, we use *K* = 256 directions and *τ* = 0.10 in our experiments. Scores are averaged across initial conditions, and summary curves plot these averages versus training epoch. We do not apply any reflection or Procrustes alignment to the point clouds prior to computing the SW2 distances.

### Task Details

#### Classification Tasks

Pre-existing labelled datasets were used for all classification tasks.

##### Breast Cancer task

The objective was to classify a breast mass as either malignant or benign using features extracted from an image of a fine needle aspirate. Thirty features were derived from ten measurements of the cell nucleus: radius, texture, perimeter, area, smoothness, compactness, concavity, number of concave points, symmetry, and fractal dimension—by computing each measurement’s mean, standard error, and the mean of its three largest values.

##### MNIST task

In this task, the goal was to classify a 28 × 28 pixel input image into one of ten categories. Each image depicted a handwritten digit, with the classes corresponding to the digits zero to nine. The dataset was accessed using the `digitTrain4DArrayData` and `digitTest4DArrayData` functions in Matlab. This is sometimes referred to as the reduced MNIST dataset, as it comprises 10,000 images instead of 70,000.

##### Fashion MNIST task

In this task, the goal was to classify a 28 × 28 pixel input image into one of ten categories. Each image depicted an item of clothing.

#### Regression Tasks

Pre-existing labelled datasets were used for all regression tasks.

##### Car Price task

The objective was to predict the price of a used Toyota car in the United Kingdom based on eight input features: registration year, transmission type, mileage, fuel type, tax, miles per gallon (MPG), and engine size.

##### Abalone Age task

This task involved predicting the age of an abalone using eight input features: sex, length, diameter perpendicular to the length, height, whole weight, shucked weight, viscera weight, and shell weight.

##### Yacht hydrodynamics task

The task was to output the residuary resistance per unit weight of displacement of yachts based on the following six input features: longitudinal position of the centre of buoyancy, prismatic coefficient, length-displacement ratio, beam-draught ratio, length-beam ratio, Froude number.

#### Dynamical Systems Tasks

For all dynamical systems tasks, the network input was the current state of the dynamical system, and the output was the next state of the system. For open-loop samples, the input was an external supervisor and for closed-loop samples, the input was the network’s previous output. All testing was performed closed-loop. The three systems considered were the Van der Pol oscillator (two-dimensional), the Lorenz system (three-dimensional, chaotic) and the Sprott S system (three-dimensional, chaotic). The true system’s dynamics were defined by the following equations:

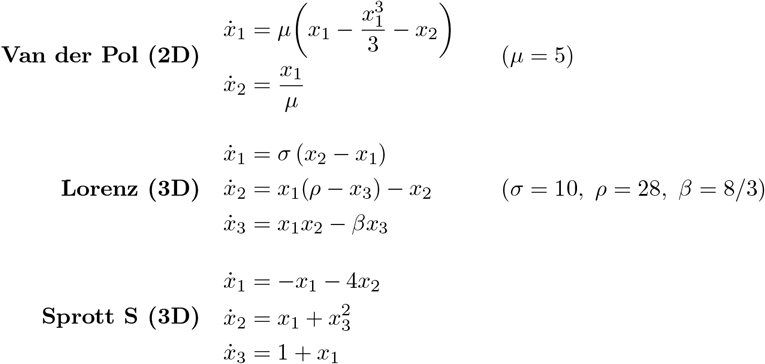

**x** = (*x*_1_, *x*_2_)^⊤^ for Van der Pol and **x** = (*x*_1_, *x*_2_, *x*_3_)^⊤^ for Lorenz and Sprott S.

## Supplementary Figures

**Supplementary Figure 1:**
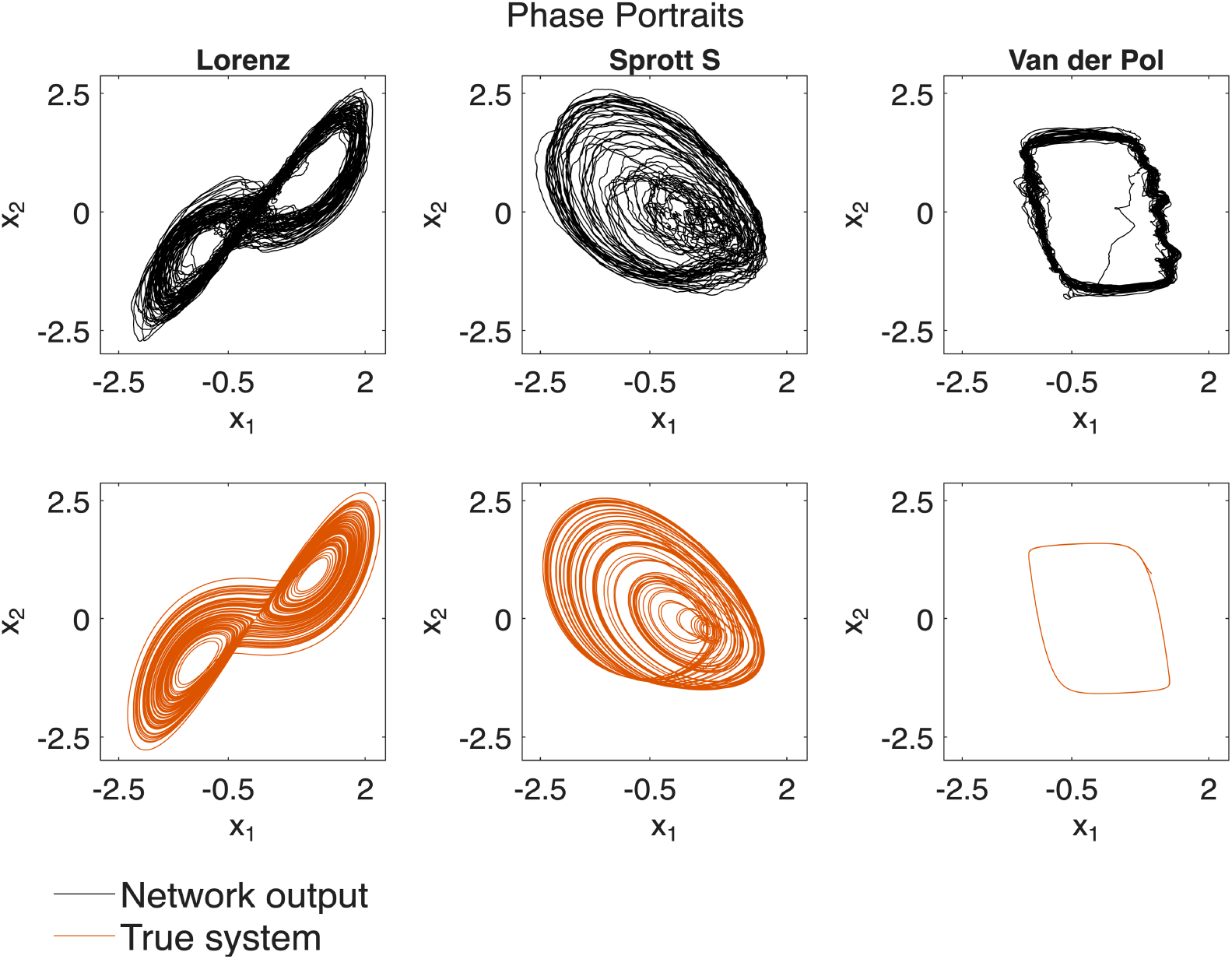
Phase portraits. The phase portraits of the network output (secondary head) and true systems for the Lorenz, Sprott S and Van der Pol systems.

**Supplementary Figure 2:**
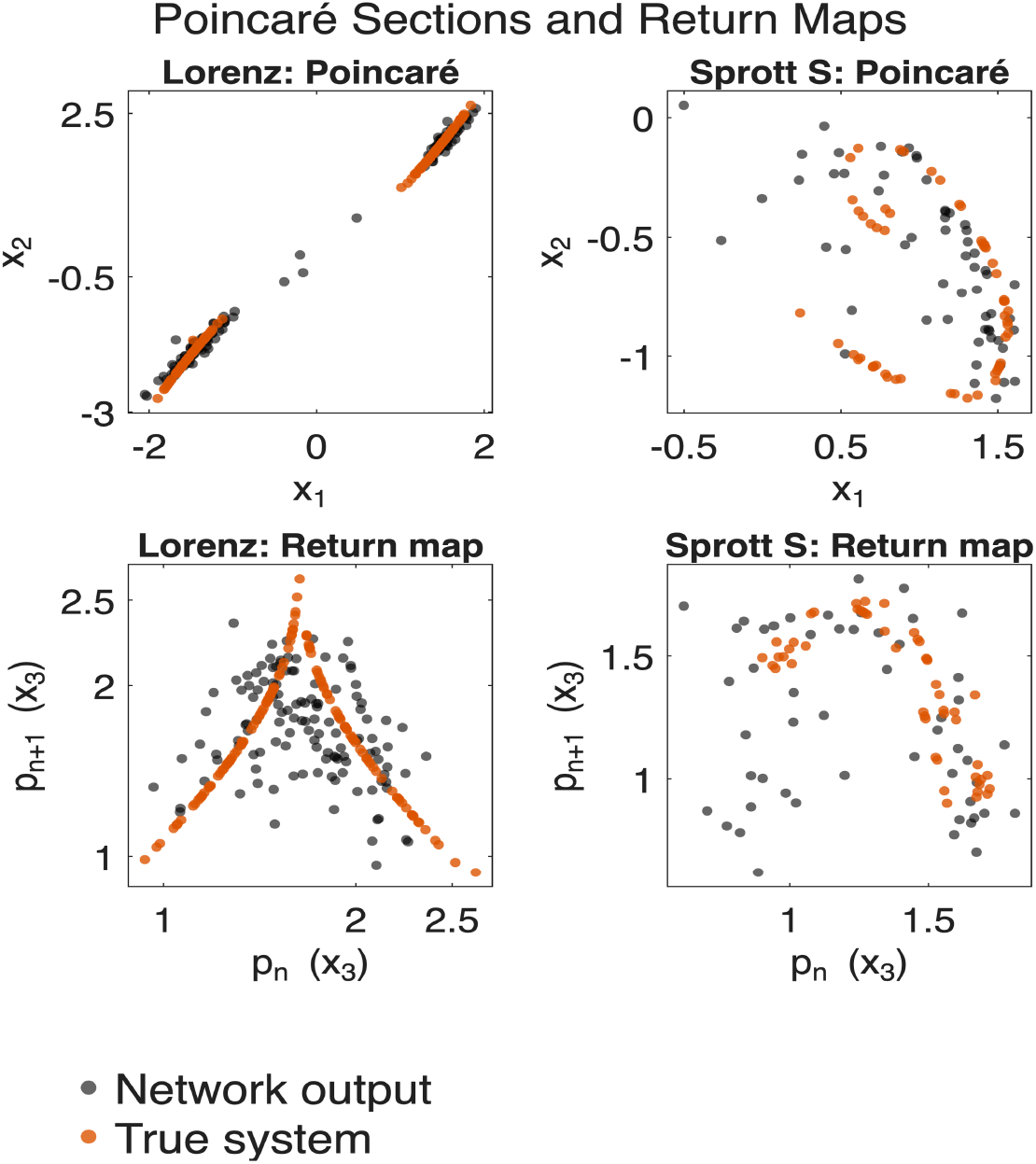
Peak return maps & Poincaré sections. The peak return maps (bottom) and Poincaré maps (top) for the closed-loop network output and true system for the Lorenz and Sprott S dynamical systems. The peak return maps plot the amplitude of the n^th^ peak versus the n+1^th^ peak for the third dimension of the system using a minimum peak prominence of one. The Poincaré maps are the zero-crossing Poincaré maps of the third dimension of the system. N.B. Return maps and Poincaré sections are not shown for the Van der Pol system because it is not a chaotic system with distinctive maps/sections.

**Supplementary Figure 3:**
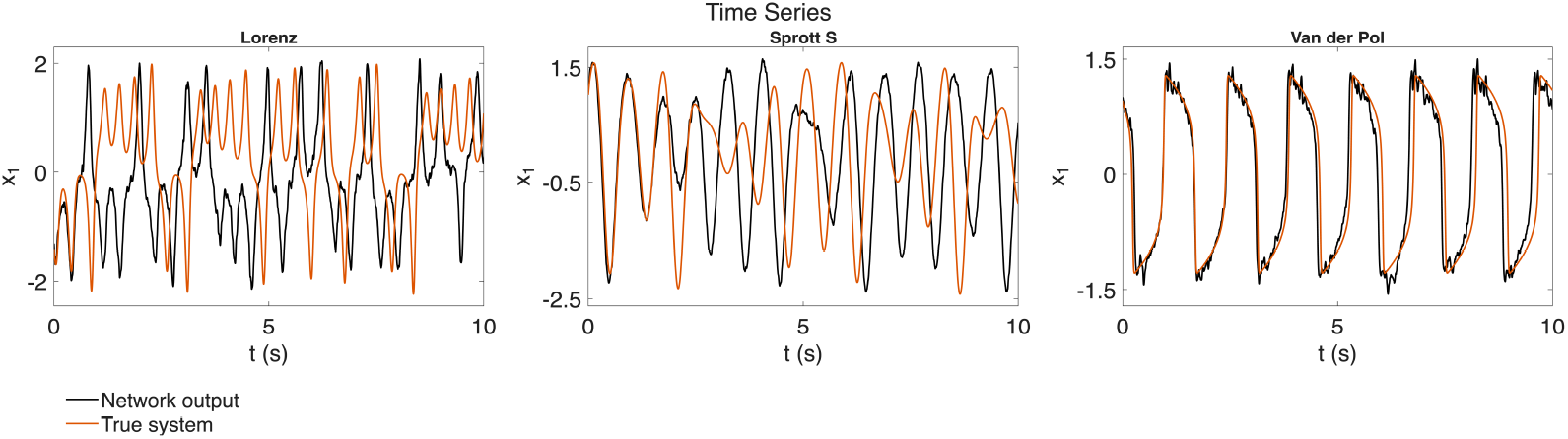
Time series. The closed-loop network output (secondary head) and true system time series for the Lorenz, Sprott S and Van der Pol systems.

